# Variation in Genome-wide NF-kappaB RELA Binding Sites upon Microbial Stimuli and Identification of a Virus Response Profile

**DOI:** 10.1101/247965

**Authors:** Lisa Borghini, Jinhua Lu, Martin Hibberd, Sonia Davila

## Abstract

NF-kB transcription factors are master regulators of the innate immune response. Activated downstream of pathogen recognition receptors, they regulate the expression of genes to help fighting infections as well as recruiting the adaptive immune system. NF-kB responds to a wide variety of signals, but the processes by which stimulus-specificity is attained remain unclear. Here, we characterized the response of one NF-kB member, RELA, to four stimuli mimicking infection in human nasopharyngeal epithelial cells. Comparing genome-wide RELA binding, we observed stimulus-specific sites, although most sites overlapped across stimuli. Specifically, the response to Poly I:C – mimicking viral dsRNA and signalling through TLR3 – induced a distinct RELA profile, binding in the vicinity of antiviral genes and correlating with corresponding gene expression. This group of binding sites was also enriched in Interferon Regulatory Factor (IRF) motifs and showed overlapping with IRFs binding sites. A novel NF-kB target, *OASL* was further validated and showed TLR3-specific activation. This work showed that some RELA DNA binding sites varied in activation response following different stimulations and that interaction with more specialized factors could help achieve this stimulus-specific activity. Our data provide a genomic view of regulated host response to different pathogen stimuli.

## Introduction

The Nuclear Factor kappa B (NF-kB) family of transcription factors (TF) is a master regulator, particularly crucial in immunity ^1^. There are five constitutively expressed NF-kB members, RELA (also called p65), RELB, REL, NFKB1 (p50) and NFKB2 (p52). They form homo or heterodimers and are kept inactive in the cytoplasm by the inhibitors of kappa B proteins (IkB). Upon stimulation, IkBs are degraded and release active NF-kB dimers that translocate into the nucleus and bind DNA to regulate gene expression. The canonical NF-kB pathway, which involves predominantly RELA:NFKB1 heterodimers, is essential in innate immunity as it regulates pro-inflammatory molecules like cytokines and chemokines as well as antimicrobials ^2^. This pathway is activated downstream of Pattern Recognition Receptors (PRRs) such as Toll like receptors (TLR) and is one of the first modes of defence against infection. TLRs are the most studied PRRs, each of them recognizes a distinct pattern derived from different human pathogens (bacteria, viruses, fungi) although they vary in their location and signalling ^3^.

NF-kB members respond to a vast array of stimuli and control the expression of many target genes but response specificity is achieved by regulation of only a subset of them, through cellular response after a particular stimulation ^4^. Stimulus-specific NF-kB activity is a complex process involving many layers of regulation ^5^ that remains unclear. The kinetic of NF-kB activation is a key factor as timing and duration of NF-kB activation varies in a stimulus-specific manner leading to variations in the regulation of genes and their kinetics of expression. A substantial amount of work has been done on characterizing the dynamics of NF-kB signalling ^6^. However, these studies focus only on one or two stimuli, generally tumour necrosis factor alpha (TNFα) and lipopolysaccharide (LPS). The formation of diverse NF-kB dimeric species following stimulation is also crucial as different dimers can regulate unique sets of target genes. NF-kB members are able to form diverse dimers by using different combinations ^7^. The separation between the canonical and non-canonical pathways activating RELA:NFKB1 or RELB:NFKB2 respectively ^8^ is a clear example of this phenomenon. Finally, NF-kB interaction with other TFs is central as well and modulates the regulation of specific genes. For instance, interaction of RELA:NFKB1 with E2F1 is necessary to fully induce a subset of pro-inflammatory cytokines following LPS stimulation ^9^.

In order to shed light on stimulus-specific NF-kB activity following PRRs stimulation, we focused on RELA’s DNA-binding function – its final role as a TF – in nasopharyngeal epithelial cells. We selected RELA due to its importance in the canonical pathway and because, unlike NFKB1, it possesses a transactivation domain that allows it to control gene transcription ^8^. We concentrated on its role in innate immunity in epithelial cells as these are the gateway to many pathogens. Following microbial recognition, epithelial cells react by producing anti-microbial molecules and pro-inflammatory factors to recruit immune cells and by interacting with cells involved in adaptive immunity. Thus they contribute greatly to the host-pathogen interactions and the resulting immune response ^10^. Detroit 562 cells were chosen due to their use as infection model to several pathogens ^11–13^. We characterized RELA activity, its DNA binding sites as well as gene expression, following stimulation with four different stimuli (Supplementary Figure S1): (1) LPS, an endotoxin present in the outer membrane of gram negative bacteria, ligand of TLR4; (2) Pam2CSK4, a chemical component mimicking bacterial peptidoglycan and recognized by TLR2; (3) Poly I:C, double stranded RNA (dsRNA) modelling viral infection through TLR3; (4) TNFα, a cytokine induced by NF-kB, which signals through the TNF receptor 1 (TNF-R1) to activate the canonical NF-kB pathway ^2^. We found differences between RELA activities upon these diverse stimuli, notably regarding its binding sites, which could explain some of its stimulus-specific activity.

## Results

### The four stimuli activate RELA and trigger the expected immune response

Detroit 562 cells were treated with LPS, TNFα, Pam2CSK4 or Poly I:C followed by extraction of nuclear protein to test RELA activation. Under resting condition, RELA was not active (low signal) but the four treatments were able to activate it (Supplementary Figure S2). Notably, the signal observed was specific as adding wild-type RELA binding sites to the reaction abolished it whereas mutated binding sites did not (Supplementary Figure S2). A time course of RELA activation over two hours was then performed and revealed that the kinetics of RELA activation varies across stimuli (Figure 1 A). Stimulation with TNFα showed the quickest response in less than 60 minutes while treatment with Poly I:C was the slowest at 90 minutes. The RELA activation peak for Pam2CSK4 was seen around 60 minutes and LPS treatment gave a more sustained activation between 60 to 90 minutes. In addition, the expression of *TNF* – considered an early NF-kB target gene following LPS treatment ^14^ – was evaluated by RT-qPCR at different time points and a similar time course of expression was observed, although it was slightly delayed as compared to that of RELA activation (Supplementary Figure S2). Taken together, these results indicate that the kinetics of NF-kB activation following the various treatments differs in a stimulus-specific manner.

**Figure 1:**
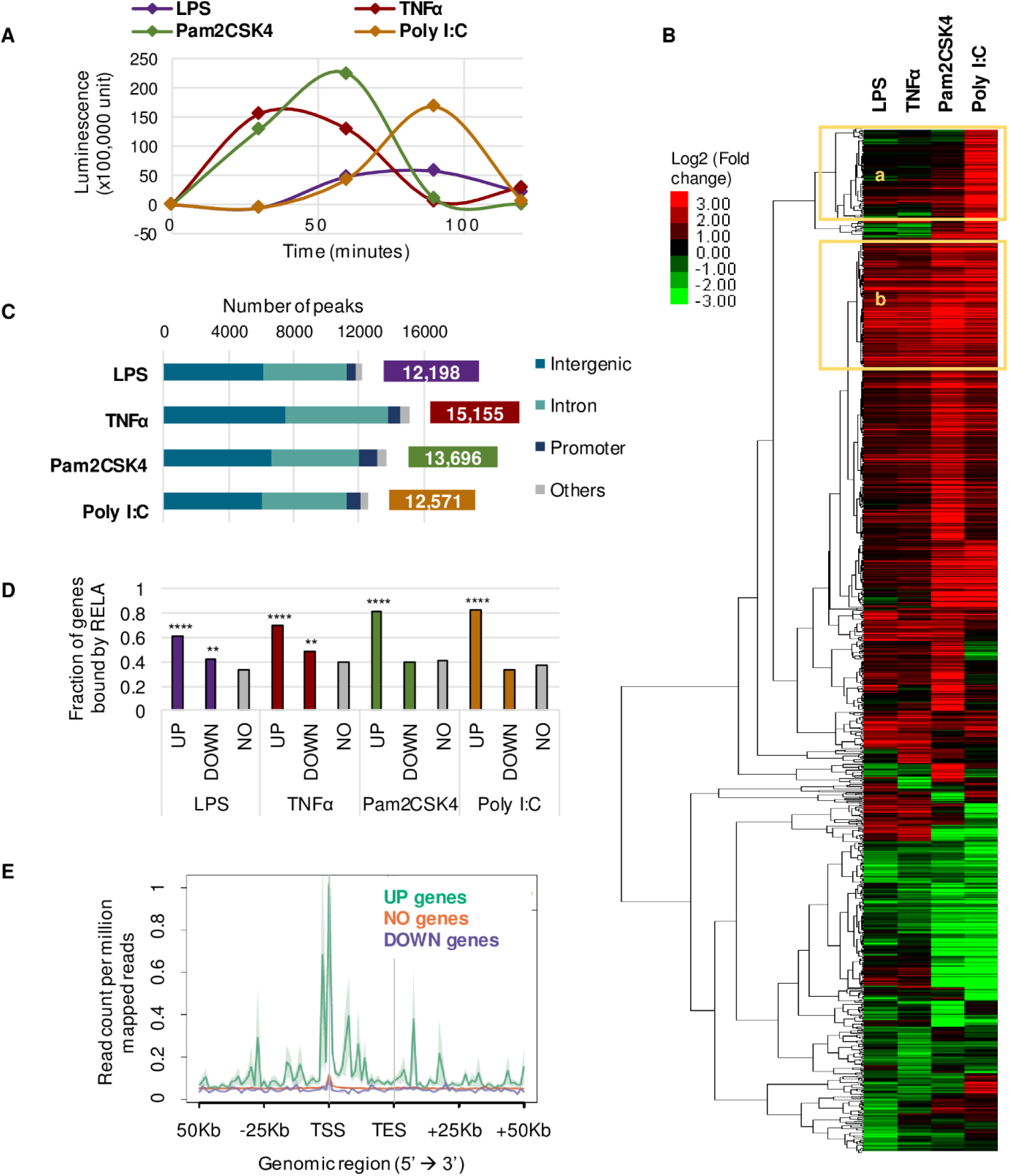
Stimulation of epithelial cells, RELA activation, DNA binding and gene expression. **A**: Time course of RELA activation. Detroit 562 cells were stimulated for two hours with LPS, TNFα, Pam2CSK4 or Poly I:C and RELA activation was determined with the NFκB p65 transcription factor assay at 30 minutes interval for 2 hours. Results are average luminescence readouts of two independent experiments. **B**: Differential gene expression in response to different stimuli. After stimulation with LPS for 100 minutes, TNFα for 70 minutes, Pam2CSK4 for 80 minutes or Poly I:C for 110 minutes, RNAs were isolated from Detroit 562 cells for RNA-seq analysis. Average Log2 fold change of biological duplicates were used for hierarchical clustering of genes differentially expressed in at least one condition. Sets of genes up-regulated in Poly I:C only (a) or all conditions (b) are highlighted. **C**: Annotation of RELA binding sites. ChIP-seq experiment was performed after treatment of Detroit 562 cells with LPS for 80 minutes, TNFα for 50 minutes, Pam2CSK4 for 60 minutes or Poly I:C for 90 minutes. The number of peaks identified for each stimulus is reported on the right. Peaks were annotated, the bar chart shows the number of peaks in different genomic features indicated. **D**: Up-regulated (UP), down-regulated (DOWN) or non-regulated (NO) genes after treatment with the four stimuli were assigned RELA ChIP-seq peaks located within 50Kb upstream and downstream of the gene body, the bar graph shows the fraction of genes assigned to at least one RELA binding site. Pearson P-values from Chi-square test against the fraction of NO genes bound by RELA are indicated, ** P<=0.01; **** P<=0.0001 **E**: RELA binding around differentially expressed genes. ChIP-seq signal upon LPS stimulation was plotted 50Kb upstream and downstream of UP, DOWN or NO genes after LPS treatment.

In order to further characterize the response to different stimuli, we evaluated differences in gene expression by RNA-seq following stimulation. We identified differentially expressed (DE) genes in each condition, there were 221 to 305 up-regulated and 66 to 246 down-regulated genes (Supplementary Figure S3). Four known NF-kB target genes – *TNF, NFKBIA, IL6* and *ICAM1* – were validated by RT-qPCR (Supplementary Figure S3). Comparison of DE genes between any two conditions displayed a relatively small overlap (Supplementary Figure S3). However, well established NF-kB targets such as *IL1B, TNFAIP2, TNFAIP3* and *CXCL8* were found among the common up-regulated genes as previously shown following LPS treatment ^9^.

We then performed differential gene expression clustering across stimuli (Figure 1 B and Supplementary Table S1). Expectedly, Gene Ontology (GO) analysis on set “b” genes – over-expressed in all conditions – revealed the enrichment of the biological processes terms “Inflammatory response” (P-value = 3.885×10^−19^) as well as “Immune response” (P-value = 3.585×10^−10^) and “I-kappaB kinase/NF-kappaB signalling” (P-value = 6.553×10^−6^) among others. In addition, the set “a” genes which consist of genes up-regulated only under Poly I:C stimulation (a ligand mimicking viral dsRNA), displayed “Defence response to virus” (P-value = 4.633×10^−17^) as top GO term (Supplementary Table S2). This indicates that the treatments with the stimuli were successful in triggering the expected immune response. While some genes were regulated to a similar extent across conditions, we detected important stimulus-specific gene expression responses as well, especially for Poly I:C stimulation.

### RELA binds DNA in a similar manner regardless of which stimulus is used for activation

To determine RELA DNA binding for each stimulus, we carried out RELA ChIP-seq experiments at the time of maximal activation in each condition. Peak calling identified 12,198 peaks under LPS, 15,155 under TNFα, 13,696 under Pam2CSK4 and 12,571 under Poly I:C (Figure 1 C). In addition, the fractions of reads inside peaks (FRIP) were calculated as a measure of noise ^15^ and were similar across conditions (Supplementary Table S3) suggesting that variation between experiments was small.

The peaks identified were annotated, proportions of the different genomic features were comparable across stimuli (Figure 1 C and Supplementary Table S4 for details) and in agreement with previous RELA ChIP-seq studies ^16,17^. The vast majority of peaks were found in intergenic regions (49.1%), as well as in introns (40.6%) while promoters contained a small portion of peaks (6.6%). We also compared the RELA sites identified in our study with known RELA binding sites and found 20.8% of them to be overlapping (4,494 out of 21,645 total peaks) with the data set downloaded from ENCODE (Supplementary Figure S4).

Additionally, GO analysis of nearby genes was performed and expectedly reported terms such as “Response to other organism” (Binomial FDR Q-value<10^−60^), “Regulation of innate immune response” (Binomial FDR Q-value<10^−32^) and “Regulation of I-kappaB kinase/NF-kappaB cascade” (Binomial FDR Q-value<10^−23^) among the most enriched (Supplementary Table S5). Motif analysis was also carried out to further validate our ChIP-seq experiment (Supplementary Table S6). As expected, NF-kB-p65 motifs were enriched in every set of peaks (-log(P-value)>2,400) much like motifs for the AP-1 complex, including Fosl2, Fra1, Jun-AP1, ATF3, BATF and AP-1 (-log(P-value)>3,000 for all), which cooperates with NF-kB ^18^.

Finally, we examined the correlation between RELA binding and gene expression by looking at RELA peaks within a gene’s regulatory region defined by the region spanning from 50Kb upstream to 50Kb downstream of the gene body. This threshold had been used previously for RELA binding and shown to be associated with up-regulation of NF-kB-dependent LPS-induced genes 9. In the case of stimulation by LPS and TNFα, up-regulated (UP) and down-regulated (DOWN) genes were significantly more bound by RELA compared to non-regulated (NO) genes while only UP genes were found to be significantly more bound by RELA in Pam2CSK4, Poly I:C conditions (Figure 1 D). In addition, for those genes with at least one RELA binding site in their regulatory region, we found that UP genes had significantly more RELA binding sites with higher p-value – used as a measure of binding strength – in their regulatory region than DOWN or NO genes (Supplementary Figure S4) showing that an increase of gene expression is more likely if multiple strong RELA binding sites are found in the proximity of a gene upon stimulation. Similarly, plotting the RELA ChIP-seq signal within and around genes showed the same trend (see LPS condition in Figure 1 E, other conditions in Supplementary Figure S4) which is in agreement with the role of RELA as a transcriptional activator ^9^, although RELA-mediated transcriptional repression has been reported as well ^19^.

Taken together, these analyses indicate that, regardless of which stimulus activates RELA, it binds DNA in a similar manner and correlates with an increase in gene expression. Furthermore, these results are in agreement with the role of RELA as a master regulator of the innate immune response and indicate successful RELA ChIP-seq experiments in all conditions.

### A minority of RELA binding sites are stimulus-specific

First, we compared the peaks identified in each condition based on their location (Figure 2 A). Surprisingly, a very large fraction of RELA binding sites were common among all stimuli (6,778 peaks out of 21,645 total peaks) and the overlap between any two conditions was very high (Supplementary Figure S5). We investigated if the non-common peaks were low confidence peaks that were therefore not called in all conditions. Comparing decreasing number of most confident peaks showed a constant decrease in the number of common peaks across stimuli (Supplementary Figure S5) supporting the existence of genuine RELA binding sites that are condition-specific.

**Figure 2:**
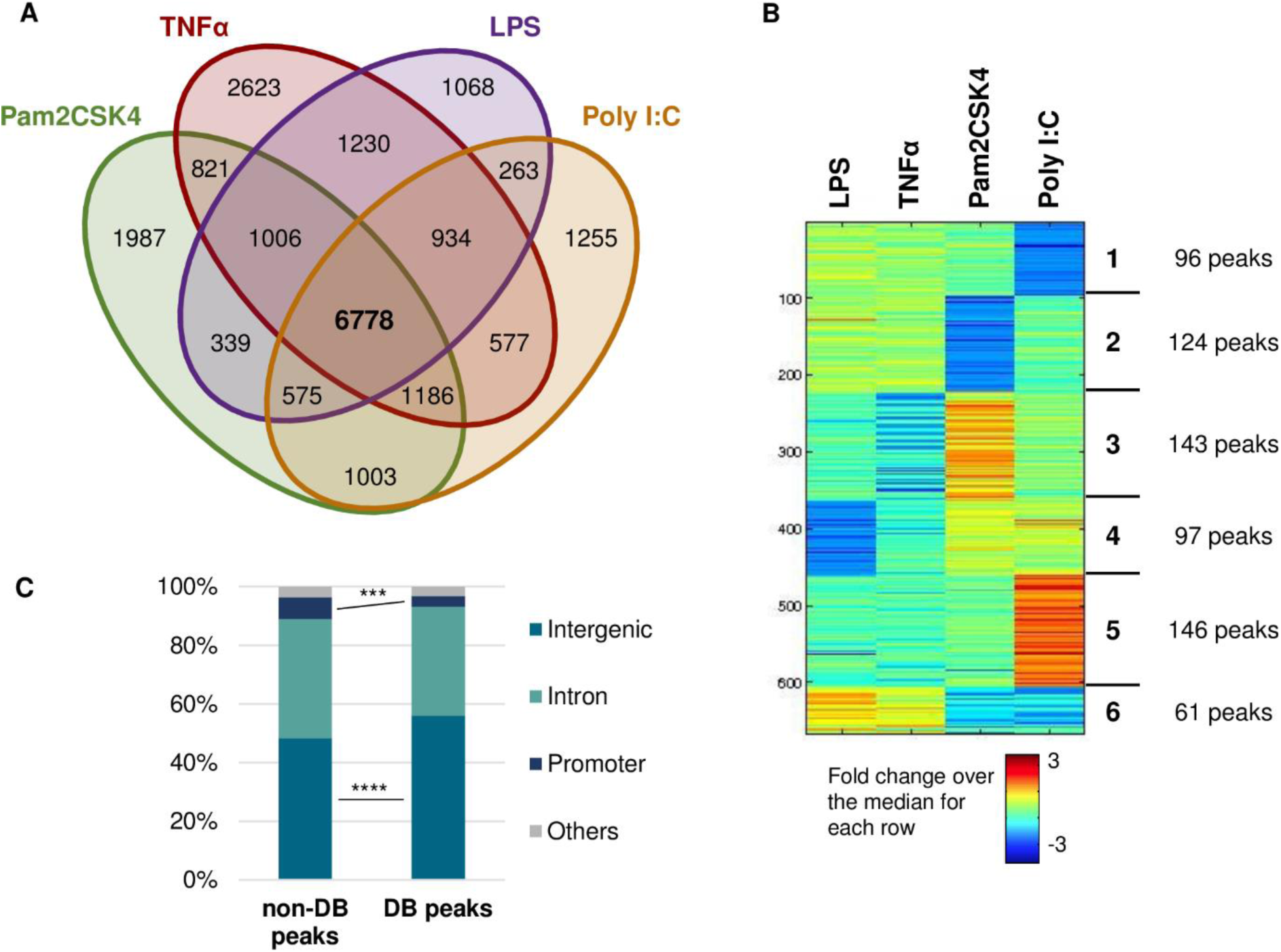
Comparison of RELA binding sites across stimuli. **A**: Comparison of RELA binding sites location. Genomic coordinates of RELA ChIP-seq peaks called upon each stimuli were compared and considered overlapping when the maximum distance between 2 peaks centers was less than 500bp. The Venn diagram shows the number of peaks overlapping across the four conditions. **B**: Differential binding analysis. The heatmap shows K-mean clustering of the ChIP-seq signal (pooled duplicates) across the four stimulations. Differentially bound regions were separated into 6 groups with the number of peaks in each group shown on the right. **C**: Annotation of Differentially bound (DB) and non-differentially bound (non-DB) peaks. The percentage of peaks in each genomic feature for both sets of peaks is reported. Pearson P-values from Chi-square test between the two sets are reported, *** P<=0.001; **** P<=0.0001.

To further compare the different sets of peaks, we performed a differential binding analysis (Figure 2 B). 667 peaks were found to be differentially bound (DB) by RELA across stimuli when a threshold of two fold change (see Methods) was used. This threshold seemed to remove most of the noise as it did not show any significant difference in terms of fraction of DB peaks between all and most confident peaks (Supplementary Figure S5). These 667 peaks were clustered into 6 groups by K-means clustering based on their binding patterns in the different conditions. While LPS and TNFα showed a similar profile, the other 2 stimuli were more distinct, particularly Poly I:C which revealed the larger group of most differentially bound peaks (Group 5 – Figure 2 B). We further characterized the DB peaks by annotating them and found that they were significantly more intergenic and significantly less promoter peaks in DB than non-DB peaks (Figure 2 C).

### The group of Poly I:C-increased peaks shows biological relevance

When we performed GO analysis on the different groups of DB peaks, Group 5 – higher RELA binding under Poly I:C stimulation – revealed enrichment for “Defence response to virus” (Hyper raw P-value = 1.64490×10^−8^) and “Regulation of TLR3 signalling pathway” (Hyper raw P-value = 6.18064×10^−7^) as top enriched biological processes terms (Figure 3 A). This matches the stimulus mimicked by this compound and shows the relevance of this particular set of peaks which we decided to study deeper. In an attempt to correlate stimulus-specific RELA binding to stimulus-specific gene expression, we assigned differentially expressed (DE) genes to DB RELA peaks located within 50Kb and looked at their expression in each condition, grouping them according to the differential binding analysis groups (Supplementary Table S7). Group 5 displayed a clear up-regulation of genes under Poly I:C stimulation (Figure 3 B), following RELA binding specificity and further confirming RELA as a transcriptional activator. Out of the 30 DE genes assigned to the peaks in this group, 18 of them were expressed at a higher level (Log2FC difference > 1) under Poly I:C than in the other conditions. Many were related to “response to virus” function such as *DD×60* or *ISG15* which were further validated by RT-qPCR (Supplementary Figure S6). Moreover, 14 of these genes were found in the set A of the differential expression clustering (Figure 1 B). They constitute potential RELA targets, specifically induced following TLR3 stimulation and suggest that some of the DB peaks identified could explain a fraction of the changes in gene expression across stimuli.

**Figure 3:**
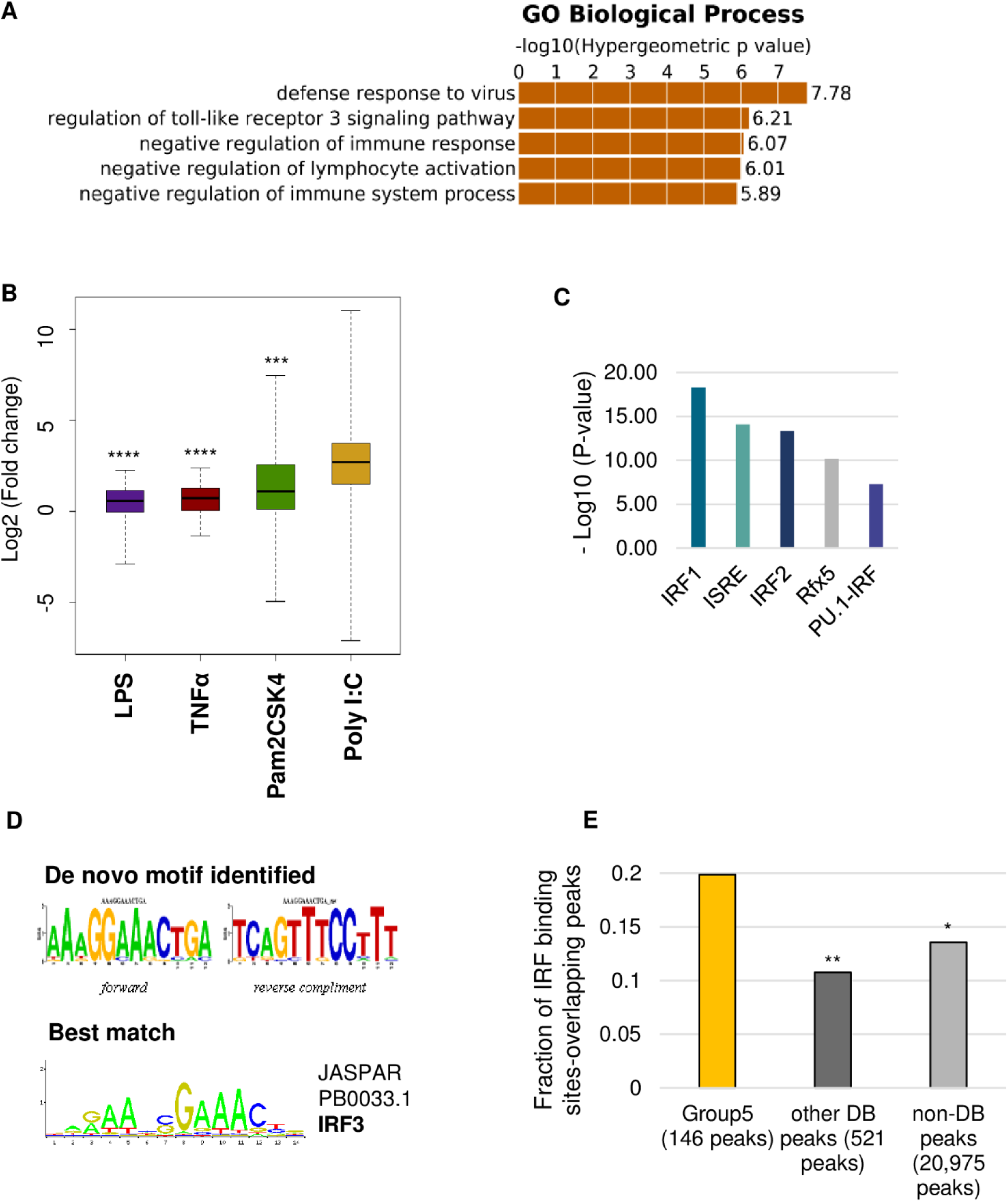
Poly I:C-increased RELA binding. **A**: Gene Ontology (GO) analysis of Group 5 peaks. Using great, genomic regions from Group 5 peaks were associated to the single nearest gene and GO analysis was performed. The top 5 Biological processes terms are reported here together with their P-value. **B**: Gene expression of genes associated with Group 5 peaks. Differentially expressed genes in at least one condition were assigned differentially bound RELA peaks located within 50Kb upstream to 50Kb downstream of the gene. The boxplot shows Log2 (Fold Change) in each condition of the genes assigned to Group 5 peaks. P-values from a paired Wilcoxon test against the gene expression under Poly I:C stimulation are indicated, *** P<=0.001; **** P<=0.0001. **C**: Motifs analysis of Group 5 peaks. Known motifs enrichment was investigated in the set of Group 5 peaks against all RELA peaks identified across stimuli. Log10 (P-values) of significantly enriched motifs are reported in the bar graph. **D**: De novo motif analysis on Group 5 peaks. Top unknown motif enriched in Group 5 peaks together with the best match from Jaspar database are represented. **E**: Overlap with IRF ChIP-seq data. Binding sites for IRFs were extracted from the ENCODE data and overlapped with the RELA peaks from Group5 (yellow), differentially bound peaks from the other groups (dark grey) or non differentially bound peaks (light grey). The fraction of RELA peaks overlapping IRF binding sites are reported, the pearson P-values from a chi-square test against the results for Group 5 peaks are indicated: ** P<=0.01; * P<=0.05.

### Interaction with Interferon Regulatory Factors could explain some of Poly I:C-specific RELA binding

In order to identify potential NF-kB partners specifically involved following TLR3 stimulation, we performed motifs enrichment analysis on Group 5 peaks using all RELA peaks identified in this study as background. Interestingly, Interferon Regulatory Factor (IRF) 1 and 2 as well as Interferon Stimulated Response Element (ISRE) motifs were highly enriched (Figure 3 C). Moreover, Rfx5 and PU.1-IRF motifs were also enriched in Group 5 peaks, to a lesser extent. De novo analysis revealed a motif that resembles the IRF3 motif (Figure 3 D) as top hit (P-value = 1×10^−15^). To further investigate the potential interaction of RELA activated via TLR3 with IRFs, we extracted IRF ChIP-seq data sets from ENCODE and overlapped these binding sites with Group 5 peaks. The latter showed a significantly higher overlap with 19.9% of IRF binding sites-overlapping peaks compared to 10.7% for the other DB peaks and 13.6% for the non-DB peaks (Figure 3 E). The ENCODE data consists of ChIP-seq for IRF1, IRF3 and IRF4, the breakdown of Group 5 RELA binding sites overlap revealed 18 common sites with IRF4, 3 with IRF3 and 17 with IRF1. A number of these overlaps were found near Poly I:C up-regulated genes (11 genes out of 18 with log2FC difference > 1 in Poly I:C compared to the other conditions) such as the ones mentioned above, *DD×60* or *ISG15* (Supplementary Figure S6). Finding enrichment for IRFs motifs and increased overlap with IRFs binding sites in this set of differentially bound peaks suggests that interacting with these factors could be a way for RELA to achieve stimulus-specificity following Poly I:C stimulation.

### OASL is a potential new NF-kB target specifically activated upon TLR3 stimulation

Among the genes associated with Group 5 peaks and showing up-regulated expression following Poly I:C stimulation only, we identified *OASL,* an antiviral gene induced by the interferon response ^20^ which is highly relevant in this case. Moreover, this gene is not known as regulated by NF-kB target as it does not appear on the list of target genes (http://www.bu.edu/nf-kb/gene-resources/target-genes/). We therefore investigated this locus as a novel TLR3-specific NF-kB target.

Our ChIP-seq data revealed a RELA binding site located around 20Kb upstream of *OASL* promoter bound more prominently under Poly I:C stimulation (Figure 4 A) that was further validated by ChIP-qPCR (Figure 4 B). Interestingly, this RELA peak overlapped an IRF1 binding site identified by the ENCODE consortium (Figure 4 A) and revealed three potential ISRE motifs in its sequence (see Supplementary methods). Gene expression analysis showed specific up-regulation following Poly I:C treatment only (Supplementary Table S1 – Ensemble ID ENSG00000135114) which was validated experimentally by RT-qPCR as well (Figure 4 C). The same expression pattern was also observed when the cells were treated with the four stimuli for the same period of time (110 minutes) showing that this change in gene expression is not due to the variable time of induction in the different conditions (Supplementary Figure S6). Furthermore, we proceeded to block NF-kB activation with the widely used inhibitor BAY 11-7082 ^21^, to investigate *OASL* regulation. BAY 11-7082 was successful in inhibiting RELA activation as well as abolishing up-regulation of known NF-kB targets (Supplementary Figure S7). In regards to *OASL,* the inhibitor also abolished binding of RELA at the potential enhancer upstream of the gene (Figure 4 D) as well as its up-regulation following Poly I:C stimulation (Figure 4 E).

**Figure 4:**
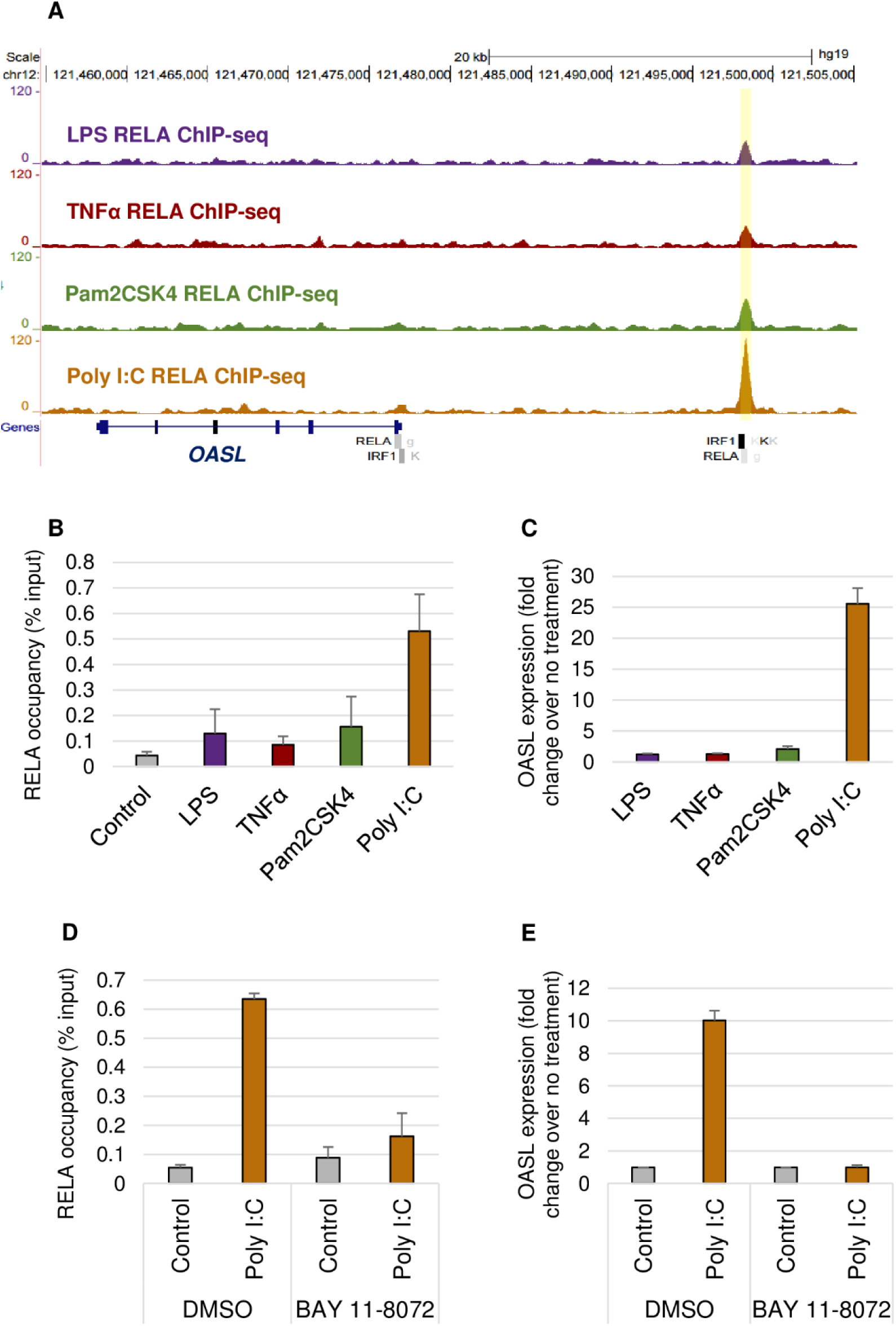
OASL locus as an example of stimulus-specific RELA target. **A**: Genome browser view of RELA ChIP-seq signal under different stimuli. A RELA binding sites with higher signal under Poly I:C stimulation is detected approximately 20Kb upstream of the *OASL* gene. **B**: Validation of the poly I:C-increased peak by ChIP-qPCR. Primers were designed to amplify the region highlighted in A (yellow). The bar chart shows the enrichment of RELA binding at this location as percent of input recovered. **C**: *OASL* expression upon five stimuli. Expression was determined by RT-qPCR and expressed as fold changes in *OASL* expression after stimulation as compared to untreated cells. **D**: Inhibition of RELA binding. Cells were pre-treated with BAY 11-8072 or DMSO followed by Poly I:C stimulation or no treatment control. Binding of RELA at the region highlighted in yellow in A was investigated by ChIP-qPCR and shown as the average input percentage recovered from the immunoprecipitation. **E**: NF-kB regulation of *OASL.* Cells were pre-treated with BAY 11-8072 or DMSO followed by Poly I:C stimulation or no treatment control and OASL expression was measured by RT-qPCR, and represented as fold change of expression over the control. Data in B, C, D and E represents the average of two independent experiments and their standard deviation as error bars.

## Discussion

When comparing the different conditions’ peaks, we were able to identify a small number of stimulus – specific binding sites. Notably, RELA binding sites with greater signal under the virus-like stimulus Poly I:C, showed particular biological relevance as they were found in proximity of genes involved in viral response and could explain up-regulation of a number of these. For instance, *OASL,* which was further validated in our system as a novel NF-kB target following TLR3 stimulation. Other potential stimulus-specific RELA target genes that emerged from our analysis will warrant further validation studies. Interestingly, IRF1 and 2, ISRE and PU1:IRF motifs were enriched in this group of DB peaks. IRFs, which bind ISRE and can interact with PU1 – to form PU1:IRF complex ^22^ -, are regulators of the interferon (IFN) response. Type I IFN response is triggered by most viruses in most cell-types, although IRF2 does not seem to be involved and IRF1 is not essential for this purpose ^23^. Nonetheless, IRF1 has been shown to interact with RELA:NFKB1 dimers at the *VCAM1* promoter to increase gene expression ^24^. Furthermore, overlap of Poly I:C-increased RELA peaks with IRF1 binding sites from ENCODE was substantial. Particularly, an IRF1 binding site – identified in K562 cells treated with IFN alpha – was found within the Poly I:C-increased RELA binding sites upstream of *OASL* validated in our study. Additionally, IRF3 motif was the best match of the *de novo* motif analysis performed on Group 5 peaks. IRF3 is particularly relevant as it is activated by TLR4 and TLR3 through the TRIF pathway ^1^. It directly interacts with RELA and generally co-activates gene expression when bound next to NF-kB ^23^. Although overlap between Poly I:C-increased RELA peaks and IRF3 binding sites from ENCODE showed limited overlap, this could be due to the low number of IRF3 binding sites in the data sets (1,722 peaks compared to 20,215 for IRF1 and 17,722 for IRF4). Moreover, the ChIP-seq data available in ENCODE come from different cell lines than the Detroit 562 cells used here which could explain the relatively low overlap. Nonetheless, enrichment for motifs of TF related to viral defence response in this particular group of RELA binding sites could provide an explanation for stimulus-specific binding. As stimulation would also trigger other pathways, some more specialized TFs would be activated and could interact and recruit RELA at specific binding sites.

Other groups of stimulus-specific RELA binding sites did not show particular distinctions. As most of the signals used were bacterial components, they may not differ enough to show any specificity, or the subtle differences between such closely related signals may not be picked up by our analyses. Furthermore, microbial stimulation would release cytokines, including TNFα, as part of the inflammatory response triggered ^3^ which would explain the similarity of the gene expression response as well as the RELA binding profile observed between TNFα and the other stimuli. In addition, time is a key factor in NF-kB signalling, for both gene-specificity ^25^ and stimulus-specificity ^26^. RNA stability and processing, especially splicing ^27^, seems to be important for controlling the timing of gene expression. Therefore, for genes expressed later, the stimulus-specific RELA binding observed here could drive stimulus-specific transcription but the corresponding change in transcript levels would only be detectable at later timings. In the same way, some of the differences seen in RELA binding could be due to the variable time points chosen for the ChIP-seq experiments across conditions. Exploring RELA DNA binding as well as gene expression at different time points following stimulation would be needed for a more complete view of NF-kB stimulus-specific activity. It is also important to mention that RELA peaks in our analysis were assigned to potential RELA targets solely based on their location (within 50 Kb upstream to 50Kb downstream of DE gene). While this approach is widely used, it could lead to false positive associations and it does not account for any long range enhancer-promoter interactions which could control gene expression. This latter point is critical as a previous study has shown that some of TNFα inducible genes expression was explained by looping of their promoters with RELA binding sites ^28^.

Comparison of the different sets of peaks revealed that a majority of the binding sites are common regardless of the stimulus used. Distinct co-factor requirements ^29^, NF-kB dimers ^17^ or chromatin changes ^30^ may be necessary for RELA to further drive gene expression. Additionally, several RELA binding sites could be required for regulation to take place ^29^. This could explain why up-regulated genes contain more RELA binding sites in their regulatory region (Supplementary Figure S4B). In addition, differentially bound peaks seemed less likely to occur at promoters compared to non-DB peaks and binding of RELA was observed at promoters of non-regulated genes suggesting the importance of enhancers. Enhancers has been shown to be especially important in cell-type specific gene expression regulation ^31^, they could play a particular role in stimulus-specific expression as well.

The time course of RELA activation showed different kinetics for the different stimuli used. The delay in the response of NF-kB to LPS stimulation compared to that of TNFα has been reported previously and related to the divergent downstream pathways of these two ligands ^32^ which could explain our results as well. TLR4 binds both adaptors MyD88 and TRIF – responsible for the early and late NF-kB activation respectively – while TLR2 uses MyD88 and TLR3 goes through TRIF exclusively ^1^. Therefore, Pam2CSK4-TLR2 would be expected to give an early NF-kB response, Poly I:C-TLR3 a later one and LPS-TLR4 a more sustained one, which is concordant with our time course experiment (Figure 1 A). Nevertheless, the previous report showed that the late NF-kB activation actually works through the induction of TNFα which then binds to its receptor and passes the signal ^32^. However, in our data, *TNF* expression following Poly I:C stimulation only increases after RELA was activated (Supplementary Figure S2C). Therefore it is unlikely that TNFα induced NF-kB activation in this case. This discordance could be due to the nature and variability of the triggered signal – induction of TNFα via TRIF adaptor pathway has only been shown in the case of LPS-TLR4 interaction – further supported by the gene expression response observed where LPS and TNFα resembled one another more than TNFα and Poly I:C (Figure 1 B). Moreover, other differences in TLR4 and TLR3 signalling, such as the use of the additional factor TRAM in TLR4 signalling, could be responsible ^33^. Another explanation of the delayed RELA activation with Poly I:C stimulation could reside in the location of the receptors as TLR3 is intracellular while the others are at the membrane. Poly I:C ligand could take longer to bind its receptor and induce signalling as it would have to go through the cellular membrane. This is not supported by the fact that the ligand DAP-containing muramyl tripeptide, which targets the intracellular NOD receptors, leads to an earlier NF-kB activation maximal at 60 minutes (Supplementary Figure S2D). Although both components may have different rates of cellular translocation and receptor binding.

When comparing the RELA binding sites identified in our study to the ones from ENCODE, we found limited overlap. The RELA sites available in the latter data set consisted in majority of the regions identified by Kasowski et al. in lymphoblastoid cell lines treated with TNFα ^34^. The difference in cell type as well as the diversity of stimuli used in our study could explain the small number of overlapping sites and stresses the importance of performing ChIP-seq experiments in different cell types to identify TF regulatory regions. In this case, NF-kB activity has been mostly studied in immune cells and therefore our study in epithelial cells stimulated with several pathogen-like stimuli notably increases the knowledge in the field.

To the best of our knowledge, this is the first study to report and compare genome-wide RELA binding under different stimuli. We characterized RELA binding sites in a nasopharyngeal epithelial cell line following stimulation with LPS, TNFα, Pam2CSK4 and Poly I:C targeting different receptors, mostly pattern recognition receptors. In addition to expanding the current repertoire of NF-kB regulatory regions, we identified stimulus-specific binding sites and potential targets of RELA. Particularly following TLR3 stimulation, which showed a distinct NF-kB response that was not well described previously. This work is relevant in regards to the innate immune response to pathogen infection, particularly airborne microbes in the nasopharynx, where the role of epithelial cells in recognizing pathogens is crucial ^35^.

## Material and Methods

### Cell culture and treatment

Detroit 562 cells were purchased from ATCC and cultured in RPMI medium (Gibco) supplemented with 10% fetal bovine serum performance (Gibco), 100U/ml penicillin and 100ug/ml streptomycin (Gibco), and 1mM sodium pyruvate (Gibco). The following ligands were used to activate NF-kB: LPS from E.coli B4:111 (Sigma) at 1μg/mL, TNFα (eBioscience) at 10ng/mL, Pam2CSK4 (Invivogen) at 1μg/mL, poly I:C (Invivogen) at 10μg/mL). The stimuli were added in the culture medium for the time indicated. Inhibition of NF-kB was achieved using BAY 11-7082 (ChemCruz) at 100uM (stock solution at 50mM in DMSO) for one hour prior and at 90uM (dilution 9/10 in the medium containing the ligand) during the treatment.

### RELA activation assay

Cells were treated with the stimuli and harvested. Cytoplasmic and nuclear proteins were extracted using the NE-PER kit (Pierce) and quantified with the Coomasie Protein quantification kit (Pierce). Ten to 15ug of nuclear proteins were used to test RELA activation using the NFκB p65 transcription factor assay kit (Pierce) according to the manufacturer’s instructions. As the luminescence readouts varies greatly from experiment to experiment, the average of 2 experiments are shown in the results (Figure 1 A) but error bars were not added. Moreover, the height of the signal across different stimuli should not be compared as they consist of individual experiments.

### ChIP-seq

Cells were treated with LPS for 80 minutes, TNFα for 50 minutes, Pam2CSK4 for 60 minutes or Poly I:C for 90 minutes. For each experiment, a “no treatment” control consisting of a change of culture medium was processed along with the treated samples. ChIP-seq was performed according to the library-on-beads protocol from Wallerman et al. ^36^ using the NEBnext DNA library prep kit (New England Biolabs) with slight modifications (see Supplementary methods for detailed protocol). Antibody against RELA was purchased from Santa Cruz Biotechnology (NF-kB p65 (c-20): sc-372). Sequencing was performed on an Illumina HiSeq sequencer (2×76bp). Biological duplicates consisting of two independent experiments were sequenced and analysed for each condition.

Sequencing reads were mapped to the human genome hg19 with BWA and quality control and filtering were then performed using samtools. Multi-aligned reads, PCR duplicates, unmapped or unmapped pair and reads with a MAPQ<20 were filtered out giving 50 to 80 million passed reads per treated sample (Supplementary Table S3). Peak calling was performed with Dfilter ^37^, peaks were identified using the irreproducibility discovery rate (IDR) method ^15^, modified to include 2 calls per sample: one against the “no treatment” sample and one against input DNA (see Supplementary Methods for details). The option-wig was added in the Dfilter command in order to generate wig files to visualize the signal in the UCSC genome browser (the signal track for one replicate per condition is shown in Figure 4 A).

Counting reads in peaks was done with the command annotatePeaks.pl (–d option) from Homer ^38^ and used to calculate the fraction of reads inside peaks (Supplementary Table S3). The same command was used for annotation of the different sets of peaks (Figure 1 C, 2 C and Supplementary Table S4). Motif analysis was also performed with Homer using findMotifsGenome.pl command. Known motif enrichment in the peak sets for each stimulus was compared against the whole genome (Supplementary Table S6) while enrichment in the different DB groups was compared against all the merged RELA peaks (Figure 3 C and 3 D). Gene ontology (GO) was carried out with GREAT, genes were associated to the peaks using the “single nearest gene” rule with a threshold of 200Kb (Supplementary Table S5 and Figure 3 A). Peak location comparison between conditions was done with the mergePeaks command from Homer using –d 500 setting and –venn option (Figure 2 A). Differential binding analysis was done after GC and read count normalization and using tag count calculated with a method described previously ^39^. Bam files from duplicates treatment and no-treatment ChIP-seq were merged with samtools and used as test and background respectively. For every site (row), normalized tag count for each condition was reported and the median calculated, fold change over the median for each stimulus was generated and used for heatmap representation. A threshold of 2 fold change over the median was chosen as it appeared not to be influenced by low confidence peaks (Supplementary Figure S5C). Clustering was then performed with K-mean clustering, dividing the data into 6 groups (Figure 2 B).

### RNA-seq

Detroit 562 cells were treated with LPS for 100 minutes, TNFα for 70 minutes, Pam2CSK4 for 80 minutes or Poly I:C for 110 minutes. Total RNAs were extracted using the RNeasy Plus Universal kit (Qiagen) and loaded on a Nano RNA Bioanalyzer chip (Agilent) for qualification and quantification. A minimum of 2 μg of RNAs with a RNA integrity number (RIN) of more than 8.5 was used for library preparation with the TruSeq Stranded mRNA Library Prep kit (Illumina) according to the manufacturer’s instructions. Samples were multiplexed and sequenced on an Illumina NextSeq (1×76bp). Biological duplicates consisting of independent experiments were sequenced for every conditions together with a “no treatment” control.

Sequencing reads were mapped with Tophat, and Cuffdiff ^40^ was used to determine differentially expressed (DE) genes(GRCh37.73 gtf file) against the control (no treatment). Genes for which the test status was not “OK” were filtered out. Then up-regulated and down-regulated genes were determined as the ones with a Log2(fold change) in both biological replicates of >1 or <−1 respectively. Clustering analysis was performed on the Log2(fold change) values with Cluster 3 ^41^ (see Supplementary methods for details). GO analysis was done with David at (<u>david-d.ncifcrf.gov</u>) ^42^ (Supplementary Table S2).

### Integration of ChIP-seq and RNA-seq data

The Rnachipintegrator (https://github.com/fls-bioinformatics-core/RnaChipIntegrator) was used to assign genes to peaks falling within 50Kb around a gene body (see Supplementary methods for details). The genes-peaks association were used to correlate RELA binding to gene expression (Figure 1 D, Supplementary Figures S4A and S4C) and to investigate expression of genes assigned to differentially bound peaks (Figure 3 B). ChIP-seq signal around and along up-, down-and non-regulated genes was plotted with the ngs.plot program ^43^ (Figure 1 E and Supplementary Figure S4C).

### Statistical analysis

Comparison of proportions/fractions in Figure 1 D and 2 C and Supplementary Figure S5C was tested using the online 2×2 Chi-square test from Vassarstats (http://vassarstats.net/odds2x2.html) for which the Pearson P-value was reported. Significance between distributions was investigated using a Wilcoxon rank-sum test (wilcox.test in R), the P-values for the boxplots in Supplementary Figure S4B and S4C were calculated using non-paired test while the ones for the boxplots in Figure 3 B were calculated with a paired test.

### Data availability

The datasets generated during and/or analysed during the current study are available in the NCBI’s Gene Expression Omnibus (GEO) repository under the following GEO Series accession numbers: GSE91018 for ChIP-seq, GSE91019 for RNA-seq and GSE91020 for the Superseries grouping these two sets of data.

## Acknowledgments

We thank Kumar Vibhor for his guidance and suggestions on the ChIP-seq analysis and his comments on the manuscript. We are thankful thank Goke Jonathan and Xingliang Liu for their help on the RNA-seq analysis. The Next Generation Sequencing Platform and the Research Pipeline Development team at the Genome Institute of Singapore made the sequencing possible as well as data processing and mapping of the reads. We also thank Tan Min Jie Alvin, Wen Fong Ooi and Kumar Vikrant for their comments on the manuscript.

### Author contributions statement

SD and LB designed the study, LB performed the experiments and the analyses, MH provided support and comments of the results, LJ contributed to the discussion. LB wrote the manuscript with the help of SD and MH.

### Competing Financial Interests

The author(s) declare no competing financial interests.

## Supplementary material

**Supplementary Methods:** Supplementary-Methods_Borghini-et-al.pdf

**Supplementary Tables:** Supplementary-Tables_Borghini-et-al.xls

**Supplementary Figures:** Supplementary-Figures_Borghini-et-al.pdf

